# Extensive impact of low-frequency variants on the phenotypic landscape at population-scale

**DOI:** 10.1101/609917

**Authors:** T. Fournier, O. Abou Saada, J. Hou, J. Peter, E. Caudal, J. Schacherer

## Abstract

Genome-wide association studies (GWAS) allows to dissect the genetic basis of complex traits at the population level^1^. However, despite the extensive number of trait-associated loci found, they often fail to explain a large part of the observed phenotypic variance^2–4^. One potential source of this discrepancy could be the preponderance of undetected low-frequency genetic variants in natural populations^5,6^. To increase the allele frequency of those variants and assess their phenotypic effects at the population level, we generated a diallel panel consisting of 3,025 hybrids, derived from pairwise crosses between a subset of natural isolates from a completely sequenced 1,011 *Saccharomyces cerevisiae* population. We examined each hybrid across a large number of growth traits, resulting in a total of 148,225 cross/trait combinations. Parental versus hybrid regression analysis showed that while most phenotypic variance is explained by additivity, a significant proportion (29%) is governed by non-additive effects. This is confirmed by the fact that a majority of complete dominance is observed in 25% of the traits. By performing GWAS on the diallel panel, we detected 1,723 significantly associated genetic variants, with 16.3% of them being low-frequency variants in the initial population. These variants, which would not be detected using classical GWAS, explain 21% of the phenotypic variance on average. Altogether, our results demonstrate that low-frequency variants should be accounted for as they contribute to a large part of the phenotypic variation observed in a population.

## Introduction

Natural populations are characterized by an astonishing phenotypic diversity. Variation observed among individuals of the same species represent a powerful raw material to have a better insight into the relation existing between genetic variants and complex traits^7^. The advances of high-throughput sequencing and phenotyping technologies greatly enhance the power of determining the genetic basis of traits in various organisms^8–11^. Dissection of the genetic mechanisms underlying natural phenotypic diversity is within easy reach by using classical mapping approaches such as linkage analysis and genome-wide association studies (GWAS)^7^. Alongside these major advances, however, it must be noted that there are some limitations. All genotype-phenotype correlation studies in humans and other model eukaryotes identified causal loci in GWAS explaining relatively little of the heritability of most complex traits^12,13^. Multiple justifications for this missing heritability have been suggested, including the presence of low-frequency variants^14–16^ as well as the low power to estimate non-additive effects^17–19^.

Among the model organisms, the budding yeast *Saccharomyces cerevisiae* is especially well suited to dissect variations observed across natural populations^20,21^. Because of their small and compact genomes, an unprecedented number of 1,011 *S. cerevisiae* natural isolates has recently been sequenced^10^, showing a high level of genetic diversity greater than that found in humans^22^. Yeast genome-wide association analyses have revealed functional Single Nucleotide Polymorphisms (SNPs), explaining a small fraction of the phenotypic variance^10^. However, these analyses highlighted the importance of the copy number variants (CNVs), which explain a larger proportion of the phenotypic variance and have greater effects on phenotypes compared to the SNPs. Nevertheless, even when CNVs and SNPs are taken together, the phenotypic variance explained is still low (around 17% on average) and consequently a large part of it is unexplained. Interestingly, much of the detected genetic polymorphisms in the 1,011 yeast genomes dataset are low-frequency variants with almost 92.7% of the polymorphic sites associated with a minor allele frequency (MAF) lower than 0.05. This trend is similar to the one observed in the human population^8,16^ and definitely raised the question of the impact of low-frequency variants on the phenotypic landscape within a population and on the missing heritability^4^. Here, we investigated the underlying genetic architecture of phenotypic variation as well as unraveling part of the missing heritability by accounting for low-frequency genetic variants at a population-wide scale and non-additive effects controlled by a single locus. For this purpose, we generated and examined a large set of traits in 3,025 hybrids, derived from pairwise crosses between a subset of natural isolates from the 1,011 *S. cerevisiae* population. This diallel crossing scheme allowed us to capture the fraction of the phenotypic variance controlled by both additive and non-additive phenomena as well as to infer the main modes of inheritance for each trait. We also took advantage of the intrinsic power of this diallel design to perform GWAS and assess the role of the low-frequency variants on complex traits.

## Results

### Diallel panel and phenotypic landscape

Based on the genomic and phenotypic data from the 1,011 *S. cerevisiae* isolates collection^10^, we selected a subset of 55 isolates that are diploid, homozygous, genetically diverse (Supplementary Fig. 1a), and coming from a broad range of ecological sources (Supplementary Fig. 1b) (*e.g.* tree exudates, *Drosophila*, fruits, fermentation processes, clinical isolates) as well as geographical origins (Europe, America, Africa and Asia) (Supplementary Fig. 1c and Supplementary Table 1). A full diallel cross panel was constructed by systematically crossing the 55 selected isolates in a pairwise manner (Supplementary Fig. 1d). In total, we generated 3,025 hybrids, representing 2,970 heterozygous hybrids with a unique parental combination and 55 homozygous hybrids. All 3,025 hybrids were viable indicating no dominant lethal interactions existing between the parental isolates. We then screened the entire set of the parental isolates and hybrids for quantification of mitotic growth abilities across 49 conditions that induce various physiological and cellular responses (Supplementary Fig. 2, Supplementary Fig. 3, Supplementary Table 2). We used growth as a proxy for fitness traits (see Methods) and this phenotyping step led to the characterization of 148,225 hybrid/trait combinations.

### Estimation of genetic variance components using the diallel panel (additive vs. non-additive)

The diallel cross design allows for the estimation of additive *vs.* non-additive genetic components contributing to each trait variation by calculating the combining abilities following Griffing’s model^23^. For each trait, the General Combining Ability (GCA) for a given parent refers to the average fitness contribution of this parental isolate across all of its corresponding hybrid combinations, whereas the Specific Combining Ability (SCA) corresponds to the residual variation unaccounted from the sum of GCAs from the parental combination. Consequently, the phenotype of a given hybrid can be formulated as µ + GCA_parent1_ + GCA_parent2_ + SCA_hybrid_, where µ is the mean fitness of the population for a given trait. We found a near perfect correlation (Pearson’s r = 0.995, p-value < 2.2e-16) between expected and observed phenotypic values, confirming the accuracy of the model used (see Methods). Using GCA and SCA values, we estimated broad-(*H^2^*) and narrow-sense (*h^2^*) heritabilities for each trait (Fig. 1). Broad-sense heritability is the fraction of phenotypic variance explained by genetic contribution. In a diallel cross, the total genetic variance is equal to the sum of GCA variance of both parents and the SCA variance in each condition. Narrow-sense heritability refers to the fraction of phenotypic variance that can be explained only by additive effects and corresponds to the variance of the GCA in each condition (Fig. 1a). The *H^2^* values for each condition range from 0.64 to 0.98, with the lowest value observed for fluconazole (1 µg.ml^−1^) and the highest for sodium meta-arsenite (2.5 mM), respectively. The additive part (*h^2^* values) ranges from 0.12 to 0.86, with the lowest value for fluconazole (1 µg.ml^−1^) and the highest for sodium meta-arsenite (2.5 mM), respectively. While broad- and narrow-sense heritabilities are variable across conditions, we can also observe that on average, most of the phenotypic variance can be explained by additive effects (mean *h^2^*=0.55). However, non-additive components contribute significantly to some traits, explaining on average one third of the phenotypic variance observed (mean *H^2^ - h^2^*=0.29) (Fig. 1a). Despite a good correlation between broad- and narrow-sense heritabilities (Pearson’s r =0.809, p-value=1.921e-12) (Fig. 1c), some traits display a larger non-additive contribution, such as in galactose (2%) or ketoconazole (10 µg/ml). Interestingly, these two conditions revealed to be mainly controlled by dominance (see below). Altogether, our results highlight the main role of additive effects in shaping complex traits at a population-scale and clearly show that this is not restricted to the single yeast cross where this trend has been first observed^24,25^. Nonetheless, non-additive effects still explain a third of the observed phenotypic variance.

**Figure 1.**
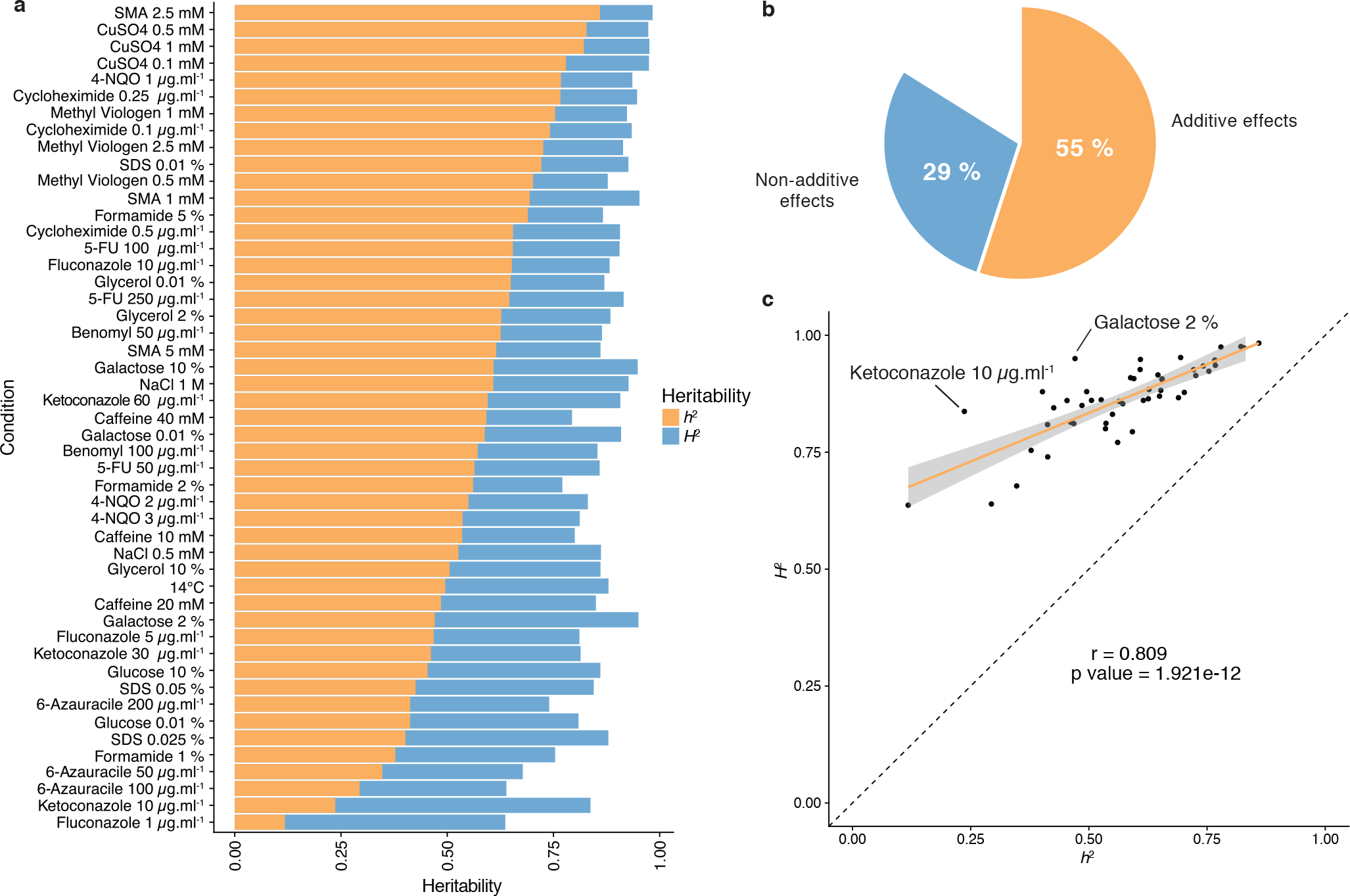
Heritability measurements. **a.** Orange bars represent the narrow-sense heritability *h*^2^ for each condition while blue bars represent broad-sense heritability *H*^2^. The difference between *H*^2^ and *h*^2^ depicts the part of variance due to non-additive effects. **b.** Overall mean additive and non-additive effects for every tested growth condition. **c.** Representation of *H*^2^ as a function of *h*^2^ showing the relative additive versus non-additive effects for each condition. Outlier conditions in terms of non-additive variance will lie further away from the linear regression line.

### Relevance of dominance for non-additive effects

To have a precise view of the non-additive components, the mode of inheritance and the relevance of dominance for genetic variance, we focused on the deviation of the hybrid phenotypes from the expected value under a full additive model. Under this model, the hybrid phenotype is expected to be equal to the mean between the two parental phenotypes, hereinafter as Mean Parental Value or Mid-Parent Value (MPV). Deviation from this MPV allows us to infer the respective mode of inheritance for each hybrid/trait combination^26^, *i.e.* additivity, partial and complete dominance towards one or the other parent and finally overdominance or underdominance (Supplementary Fig. 4, see Methods). Only 17.4% of all hybrid/trait combinations showed enough phenotypic separation between the parents and the corresponding hybrid, allowing the complete partitioning in the seven above-mentioned modes of inheritance. For the 82.6% remaining cases, only a separation of overdominance and underdominance can be achieved (Fig. 2a). Interestingly, these events are not as rare as previously described^27^, with 11.6% of overdominance and 10.1% of underdominance (Fig. 2b). When a clear separation is possible (Fig. 2c), one third of the trait/cross combinations detected are purely additive whereas the rest displays a deviation towards one of the two parents, with no bias (Fig. 2c). When looking at the inheritance mode in each condition, most of the studied traits (33 out of 49) showed a prevalence of additive effects. However, 17 of them are not predominantly additive throughout the population. Indeed, a total of 12 traits were detected as mostly dominant with 4 cases of best parent dominance, including galactose (2%) and ketoconazole (10 µg.ml^−1^), and 8 of worst parent dominance. The remaining 5 conditions display a majority of partial dominance (Fig. 2d). These results confirm the importance of additivity in the global architecture of traits. But more importantly, these results clearly demonstrate the major role of dominance as a driver for non-additive effects. Nevertheless, the presence of conditions with a high proportion of partial dominance combined with the cases of over and underdominance may indicate a strong impact of epistasis on phenotypic variation.

**Figure 2.**
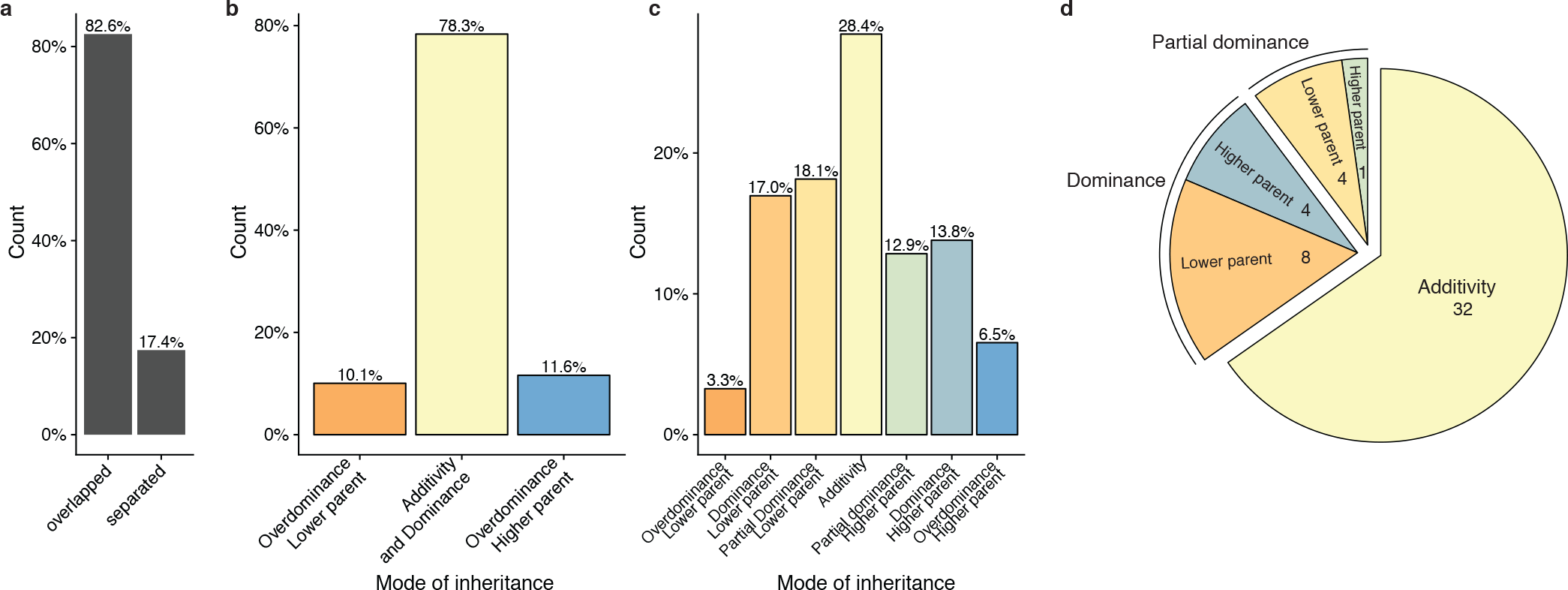
Mode of inheritance. **a.** Percentage of parental phenotypes separated from each other for which a complete partition of different inheritance modes can be achieved. **b.** Inheritance modes for every cross and condition where no separation can be achieved between the two homozygous parents. **c.** Inheritance modes for every cross and condition where a clear phenotypic separation can be achieved between the two homozygous parents. **d.** The number of conditions in each main inheritance mode.

### Diallel design allows mapping of low frequency variants in the population using GWAS

Next, we explored the contribution of low-frequency genetic variants (MAF < 0.05) to the observed phenotypic variation in our population. Genetic variants considered by GWAS need to have a relatively high frequency in the population to be detectable, usually over 0.05 for relatively small datasets^1^. Consequently, low-frequency variants are evicted from classical GWAS. However, the diallel crossing scheme stands as a powerful design to assess the phenotypic impact of low-frequency variants present in the initial population as each parental genome is presented several times, creating haplotype mixing across the matrix and preserving the detection power in GWAS.

To avoid issue due to population structure, we selected a subset of hybrids coming from 34 unrelated isolates in the original panel to perform GWAS (see Methods, Supplementary Table 1). By combining known parental genomes, we constructed *in silico* 595 hybrid genotypes matching one half matrix of the diallel plus the 34 homozygous diploids. We built a matrix of genetic variants for this panel and filtered SNPs to only retain biallelic variants with no missing calls. In addition, due to the small number of unique parental genotypes, extensive long-distance linkage disequilibrium was also removed (see Methods), leaving a total of 31,632 polymorphic sites in the diallel population. Overall, 3.8% (a total of 1,180 SNPs) had a MAF lower than 0.05 in the initial population of the 1,011 *S. cerevisiae* isolates but surpass this threshold in the diallel panel, going up to a MAF of 0.32 (Fig. 3a-b).

**Figure 3.**
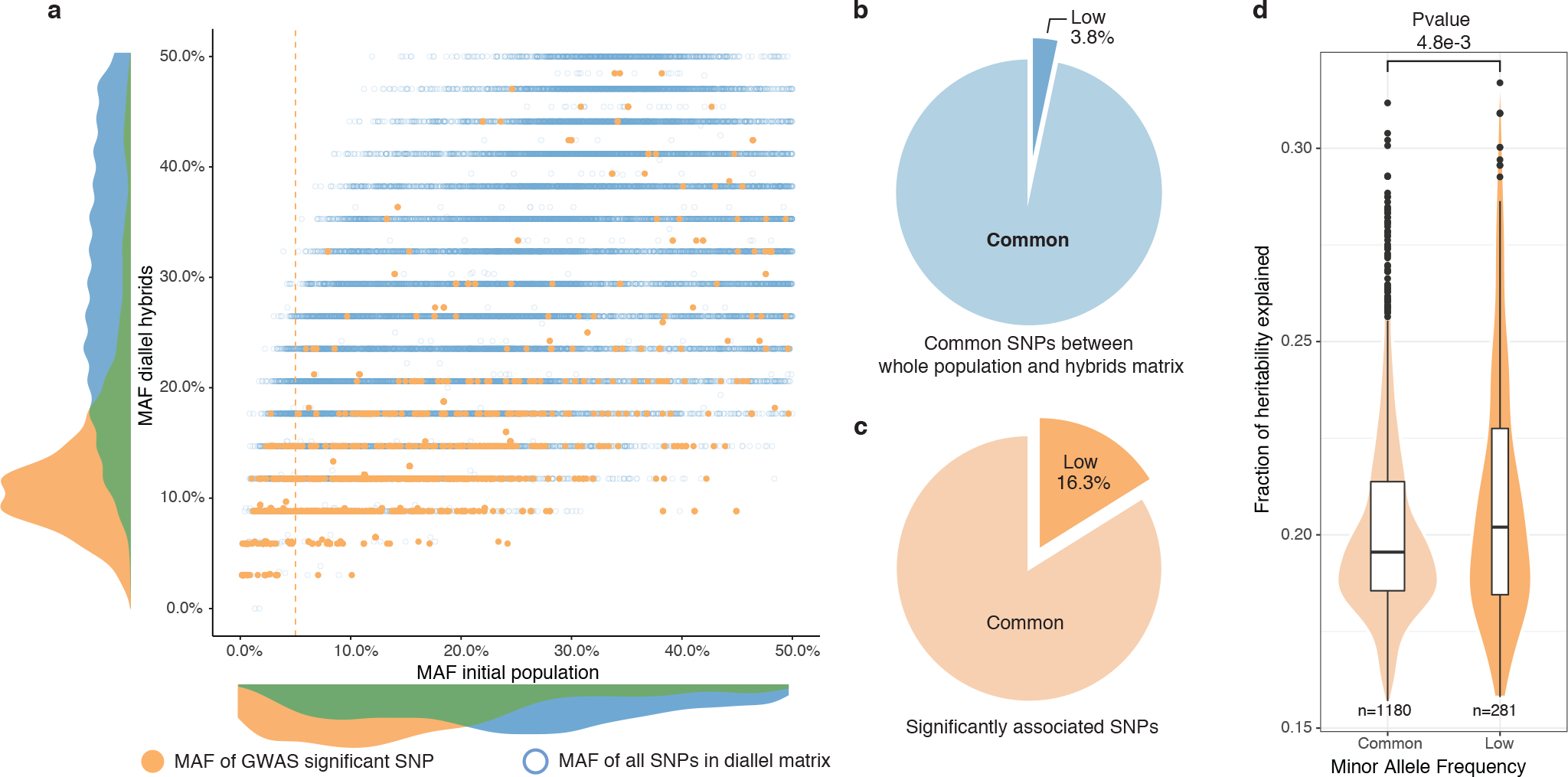
Rare and low-frequency variants detection. **a.** Comparison of MAF for each SNP between the whole population (1,011 strains) and the hybrid diallel matrix used for GWAS. Hollow blue circles represent the MAF of all SNPs common to the initial population and the diallel hybrids (31,632). Full orange circles show the MAF of significantly associated SNPs. Vertical orange line shows the 5% MAF threshold. **b.** Proportion of SNPs with a MAF below 0.05. **c.** Proportion of significantly associated SNPs with a MAF below 0.05. **d.** Fraction of heritability explained for common and low-frequency variants. P-value calculated using a two-sided Mann-Whitney-Wilcoxon test.

To map additive as well as non-additive variants impacting phenotypic variation, we performed GWA using two different models^28^ (see Methods). We used a classical additive model, encoding for SNPs where linear relationship between trait and genotype is searched, *i.e.* every locus has a different encoding for each genotype. To account for non-additive inheritance, we also used an overdominant model, which only considers differences between heterozygous and homozygous thus revealing overdominant and dominant effects. For each of these two models, we performed mixed-model association analysis of the 49 growth traits with FaST-LMM^29^. Overall, GWAS revealed 1,723 significantly associated SNPs (Supplementary Table 3) by detecting from 2 to 103 significant SNPs by condition, with an average of 39 SNPs per trait. Minor allele frequencies of the significantly associated SNPs were determined in the 1,011 sequenced genomes, from which the diallel parents were selected (Fig. 3). Interestingly, 16.3% of the significant SNPs (281 in total) correspond to low-frequency variants (MAF<0.05), with 19.5% of them (55 SNPs) being rare variants (MAF<0.01). This trend is the same and maintained for both models, with 19.3% and 15.2% of low-frequency variants for the additive and overdominant models. Because of the scheme used, it is important to note that it is possible to increase the MAF of low-frequency variants at a detectable threshold in the diallel panel and to query their effects but it is still difficult for truly rare variants (MAF<0.01), probably leading to an underestimation. However, these results clearly show that low-frequency variants indeed play a significant part in the phenotypic variance at the population-scale. We then estimated the contribution of the significant variants to total phenotypic variation (see Methods) and found that detected SNPs could explain 15% to 32% of the variance, with a median of 20% (Fig. 3b). When looking at the variance explained by each variant over their respective allele frequency, it is noteworthy that low-frequency variants explain a slightly but significantly higher proportion of the phenotypic variation (median of 20.2%) than the common SNPs (median of 19.6%) (Fig. 3b). In addition, the variance explained by the associated rare variants is also higher on average than the rest of the detected SNPs (Supplementary Fig. 5). It is noteworthy that this trend is robust and conserved across the two used encoding models, accounting for additive and overdominant effects (Supplementary Fig. 5).

To gain insight into the biological relevance of the set of associated SNPs, we first looked at the distribution across the genome and we found that 62.5% of them are in coding regions (with coding regions representing a total of 72.9% of the *S. cerevisiae* genome) (Supplementary Fig. 6a) and all these SNPs are distributed over a set of 546 genes. Over the last decade, an impressive number of quantitative trait locus (QTL) mapping experiments were performed on a myriad of phenotypes in yeast leading to the identification of 178 quantitative trait genes (QTG)^30^ and we found that 27 of the genes we detected are part of this list (Supplementary Fig. 6b). In addition, 23 associated genes were also found as overlapping with a recent large-scale linkage mapping survey in yeast^31^ (Supplementary Fig. 6b). We then asked whether the associated genes were enriched for specific gene ontology (GO) categories (Supplementary Table 4). This analysis revealed an enrichment (p-value= 5.39×10^−5^) in genes involved in “response to stimulus” and “response to stress”, which is in line with the different tested conditions leading to various physiological and cellular responses.

### *SGD1* and the mapping of a low frequency variant

Finally, we focused on one of the most strongly associated genetic variant out of the 281 low-frequency variants significantly associated with a phenotype. The chosen variant consists of two adjacent SNPs in the *SGD1* gene and has been detected in 6-azauracile (100 µg.ml^−1^) with a p-value of 2.75e-8 with the overdominant encoding and 6.26e-5 with the additive encoding. Their MAF in the initial population is only 2.5% and goes up to 9% in the diallel panel with three genetically distant strains carrying it (Fig. 6c). The SNPs are in the coding sequence of *SGD1*, an essential gene encoding a nuclear protein. The minor allele (AA) induces a synonymous change (TT**G** (Leu) → TT**A** (Leu)) for the first position and a non-synonymous mutation (**G**AA (Glu)→ **A**AA (Lys)) for the second position (Supplementary Fig. 7a). The phenotypic advantage conferred by this allele can be observed with significant differences between the homozygous for the minor allele, heterozygous and homozygous for the major allele (Supplementary Fig. 7b). To functionally validate the phenotypic effect of this low-frequency variant, CRISPR-Cas9 genome-editing was used in the three strains carrying the minor allele (AA) in order to switch it to the major allele (GG) and assess its phenotypic impact. Both mating types have been assessed for each strain. When phenotyping the wildtype strains containing the minor allele and the mutated strains with the major allele, we could see that the minor allele confers a phenotypic advantage of 0.2 growth ratio compared to the major allele (Supplementary Fig. 7c) therefore validating the important phenotypic impact of this low-frequency variant. However, no assumptions can be made regarding the exact effect of this allele at the protein-level because no precise characterization has ever been carried out on Sgd1p and no particular domain has been highlighted.

## Conclusion

Understanding the source of the missing heritability is essential to precisely address and dissect the genetic architecture of complex traits. The contribution of rare and low-frequency variants to traits is largely unexplored. In humans, these genetic variants are widespread but only few of them have been associated with some specific traits and diseases^16^. Recently, it has been shown that the missing heritability of height and body mass index is accounted for by rare variants^32^. We also recently found in yeast that most of the QTNs (Quantitative Trait Nucleotides) previously identified by linkage mapping were at low allele frequency in the 1,011 *S. cerevisiae* population^10,33,34^. This observation was corroborated by additional mapped loci via linkage mapping and analyses^31^. It also raised the question of whether these rare and large effect size alleles discovered in specific crosses are really relevant to the variation across most of the population. Here, we quantified the contribution of low-frequency variants across a large number of traits and found that among all the detected genetic variants by GWAS on a diallel panel, 16.3% of them have a low-frequency in the initial population and explain a significant part of the phenotypic variance (21% on average). This particular diallel design also presents an intrinsic power to evaluate the additive vs. non-additive genetic components contributing to the phenotypic variation. We assessed the effect of intra-locus dominance on the non-additive genetic component and showed that dominance at the single locus level contribute to the phenotypic variation observed. However, other more complicated inter-loci interactions may still be involved. Altogether, these results have major implications for our understanding of the genetic architecture of traits in the context of the unexplained heritability.

## Methods

### Construction of the diallel panel

#### Selected *Saccharomyces cerevisiae* isolates

Out of the collection of 1,011 strains^10^, a total of 53 natural isolates were carefully selected to be representative of the *Saccharomyces cerevisiae* species. We selected isolates from a broad ecological origins and we prioritized for strains that are diploid, homozygous, euploid and genetically as diverse as possible, *i.e.* up to 1% of sequence divergence. All the isolate details, including ecological and geographical origins, are listed in Supplementary table 1. In addition to these 53 isolates, we included two laboratory strains, namely ∑1278b and the reference S288c strain.

#### Generation of stable haploids

For each selected parental strain, stable haploid strains were obtained by deleting the *HO* locus. The *HO* deletions were performed using PCR fragments containing drug resistance markers flanked by homology regions up and down stream of the *HO* locus, using standard yeast transformation method (ref). Two resistance cassettes, *KanMX* and *ClonNAT*, were used for *MAT*a and *MAT* haploids, respectively. The mating-type (*MAT*a and *MAT* of antibiotic-resistant clones was determined using testers of well-known mating type. For each genetic background, we selected a *MAT*a and *MAT* clone that are resistant to G418 or nourseothricin, respectively.

Phenotyping of the parental haploid strains was performed to check for mating type specific fitness effects. All *MAT*a and *MAT* parental strains were tested on all 49 growth conditions (see below) using the same procedure as the phenotyping assay of the hybrid matrix. The overall correlation between the *MAT*a and *MAT* parental strains was 0.967 (Pearson, p-value < 1e-324), with an average correlation per strain of 0.976 across different conditions (Supplementary Fig. 8). No significant mating type specificity was identified.

#### Diallel scheme

Parental strains were arrayed and pregrown in liquid YPD (1% yeast extract, 2% peptone and 2% glucose) overnight. Mating was performed with ROTOR™ (Singer Instruments) by pinning and mixing *MAT*a over *MAT* parental strains on solid YPD. The parental strains, *i.e.* 55 *MAT*a *HO::∆KanMX* and 55 *MAT HO::∆ClonNAT* strains were arrayed and mated in a pairwise manner on YPD for 24 hours at 30°C. The mating mixtures were replicated on YPD supplemented with G418 (200 µg.ml^−1^) and nourseothricin (100 µg.ml^−1^) for double selection of hybrid individuals. After 24 hours, plates were replicated again on the same media to eliminate potential residuals of non-hybrids cells. In total, we generated 3,025 hybrids, representing 2,970 heterozygous hybrids with a unique parental combination and 55 homozygous hybrids.

### High-throughput phenotyping and growth quantification

Quantitative phenotyping was performed using endpoint colony growth on solid media. Strains were pregrown in liquid YPD medium and pinned onto a solid SC (Yeast Nitrogen Base with ammonium sulfate 6.7 g.l^−1^, amino acid mixture 2 g.l^−1^, agar 20 g.l^−1^, glucose 20 g.l^−1^) matrix plate to a 1,536 density format using the replicating ROTOR™ robot (Singer Instruments). Two replicates of each parental strain were present on every plate and six replicates were present for each hybrid. The resulting matrix plates were incubated overnight to allow sufficient growth, which were then replicated onto 49 media conditions, plus SC as a pinning control (Supplementary Fig. 2, Supplementary Table 2). The selected conditions impact a broad range of cellular responses, and multiple concentrations were tested for each compound (Supplementary Fig. 3). Most tested conditions displayed distinctive phenotypic patterns, suggesting different genetic basis for each of them (Supplementary Fig. 3). The plates were incubated for 24 hours at 30°C (except for 14°C phenotyping) and were scanned with a resolution of 600 dpi at 16-bit grayscale. Quantification of the colony size was performed using the R package Gitter^35^ and the fitness of each strain on the corresponding condition was measured by calculating the normalized growth ratio between the colony size on a condition and the colony size on SC. As each hybrid is present in six replicates, the value considered for its phenotype is the median of all its replicates, thus smoothing the effects of pinning defect or contamination. This phenotyping step led to the determination of 148,225 hybrid/trait combinations.

### Diallel combining abilities and heritabilities

Combining ability values were calculated using half diallel with unique parental combinations, excluding homozygous hybrids from identical parental strains. For each hybrid individual, the fitness value is expressed using Griffing’s model^23^:

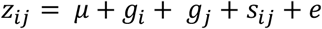

Where *z*_*ij*_ is the fitness value of the hybrid resulting from the combination of *i^th^* and *j^th^* parental strains, *μ* is the mean population fitness, *g*_*i*_ and *g*_*j*_ are the general combining ability for the i^th^ and j^th^ parental strains, *s*_*ij*_ is the specific combining ability associated with the *i* × *j* hybrid, and *e* is the error term (*i* = 1…*N*, *j* = 1…*N*, *N* = 55). General combining ability for the *i^th^* parent is calculated as:

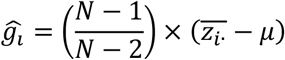

Where *N* is the total number of parental types, 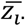 is the mean fitness value of all half sibling hybrids involving the *i*^*th*^ parent, and *μ* is the population mean. The error term associated with *g*_*i*_ is:

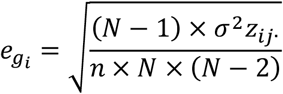

Where *N* is the total number of parental types, *n* is the number of replicates for the *i × j* hybrid, and σ^2^*z*_*ij*_. is the variance of fitness values from a full-sib family involving the *i^th^* and *j^th^* parents, which is expressed as:

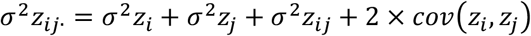

Specific combining ability for the *i* × *j* hybrid combination therefore:

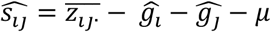

The error term associated with 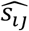 is:

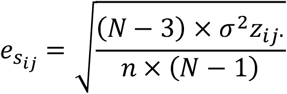

Using combining ability estimates, broad- and narrow-sense heritabilities can be calculated. Narrow sense heritability (*h*^2^) accounts for the part of phenotypic variance explained only by additive variance, expressed as the additive variance 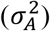 over the total phenotypic variance observed 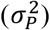:

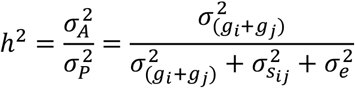

Where 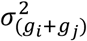 is the sum of GCA variances, 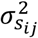 is the SCA variance and 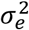 is the variance due to measurement error, which is expressed as:

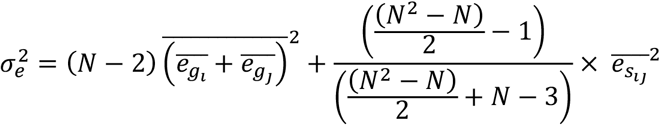

On the other hand, broad-sense heritability (*H*^2^) depicts the part of the phenotypic variance explained by the total genetic variance 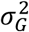:

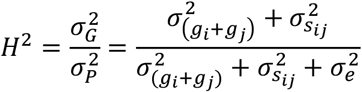

Phenotypic variance explained by non-additive variance is therefore equal to the difference between *H*^2^ and *h^2^*. All calculations were performed in R using custom scripts.

### Calculation of mid-parent values and classification of mode of inheritance

Mid-Parent Value (MPV) is expressed as the mean fitness value of both diploid homozygous parental phenotypes:

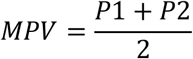

Comparing the hybrid phenotypic value (*Hyb*) to its respective parents’ allows us to infer the mode of inheritance for each hybrid/trait combination (Supplementary Fig. 4). To have a robust classification, confidence intervals for each class are based on standard deviation of hybrid (6 replicates) and parents (54 replicates). *P2* is the phenotypic value of the fittest parent while *P1* is the phenotypic value of the least fit parent.

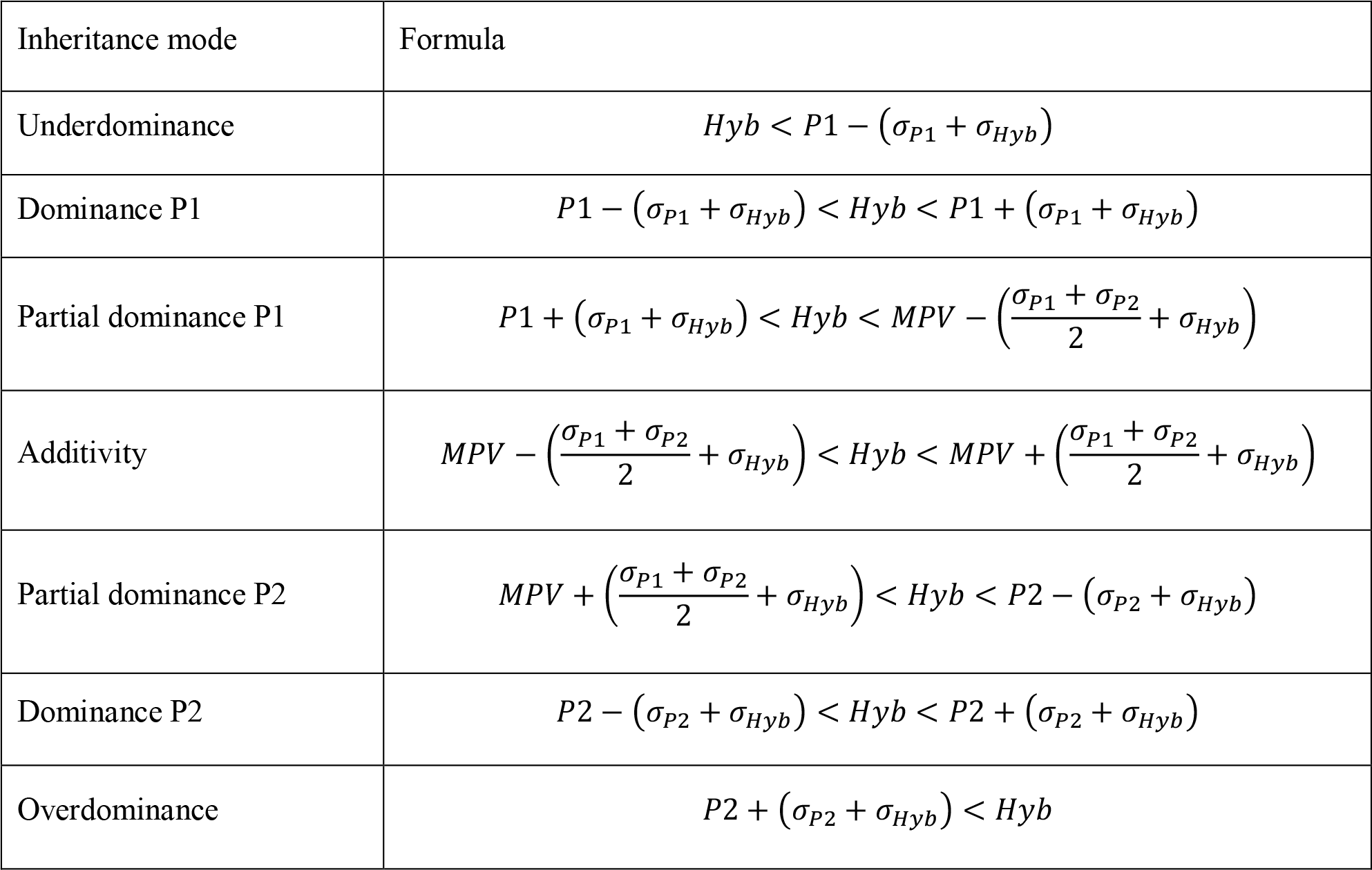

When a clear separation is possible between the two parental phenotypic values *P*1 + σ_*P*1_ < *P*2 − σ_*P*2_

The full decomposition in the 7 above mentioned categories is possible (Supplementary Fig. 4a). However, in most of the cases, the two parental phenotypic values are not separated enough to achieve this but it is still possible to distinguish between overdominance and underdominance (Supplementary Fig. 4b, Fig 2a). All calculations were performed in R using custom scripts.

### Genome-wide association studies on the diallel panel

Whole genome sequences for the parental strains were obtained from the 1002 yeast genome project^10^. Sequencing was performed by Illumina Hiseq 2000 with 102 bases read length. Reads were then mapped to S288c reference genome using bwa (v0.7.4-r385)^36^. Local realignment around indels and variant calling has been performed with GATK (v3.3-0)^37^. The genotypes of the F1 hybrids were constructed *in silico* using 34 parental genome sequences. We retained only the biallelic polymorphic sites, resulting in a matrix containing 295,346 polymorphic sites encoded using the “recode12” function in PLINK^38^. Those genotypes correspond to a half-matrix of pairwise crosses with unique parental combinations, including the diagonal, *i.e.* the 34 homozygous parental genotypes. For each cross, we combined the genotypes of both parents to generate the hybrid diploid genome. As a result, heterozygous sites correspond to sites for which the two parents had different allelic versions. We removed long-range linkage disequilibrium sites in the diallel matrix due to the low number of founder parental genotypes by removing haplotype blocks that are shared more than twice across the population, resulting in a final dataset containing 31,632 polymorphic sites.

We performed GWA analyses with different encodings^28^. In the additive model, the genotypes of the F1 progeny were simply the concatenation of the genotypes from the parents. As homozygous parental alleles were encoded as 1 or 2, the possible alleles for each site in the F1 genotype were “11” and “22” for homozygous sites and “12” for heterozygous sites. We also used an overdominant genotype encoding, where both the homozygous minor and homozygous major alleles are encoded as “11” and the heterozygous genotype is encoded as “22”.

Mixed-model association analysis was performed using the FaST-LMM python library version 0.2.32 (https://github.com/MicrosoftGenomics/FaST-LMM)^39^. We used the normalized phenotypes by replacing the observed value by the corresponding quantile from a standard normal distribution, as FaST-LMM expects normally distributed phenotypes. The command used for association testing was the following: single_snp(bedFiles, pheno_fn, count_A1=True), where bedFiles is the path to the PLINK formatted SNP data and pheno_fn is the PLINK formatted phenotype file. By default, for each SNP tested, this method excludes the chromosome in which the SNP is found from the analysis in order to avoid proximal contamination. Fast-LMM also computes for each SNP the fraction of heritability explained. The mixed model adds a polygenic term to the standard linear regression designed to circumvent the effects of relatedness and population stratification.

We estimated a trait-specific p-value threshold for each condition by permuting phenotypic values between individuals 100 times. The significance threshold was the 5% quantile (the 5th lowest p-value from the permutations). With that method, variants passing this threshold will have a 5% family-wise error rate. Taken together, GWA revealed 1,723 significantly associated SNPs (Supplementary table 3), with 1,273 and 450 SNPs for overdominant and additive model, respectively.

### Gene ontology analysis

GO term enrichment was performed using SGD GO Term Finder (https://www.yeastgenome.org/goTermFinder) with the 546 unique genes containing significantly associated SNPs. Significant enrichment is considered under “Process” ontology with a p-value cutoff of 0.05.

### CRISPR-Cas9 allele editing

pAEF5 plasmid containing Cas9 endonuclease and the guide RNA targeting *SGD1* was co-transformed with the repair fragment of 100 nucleotides containing the desired allele. Transformed cells were then plated on YPD supplemented with 200 µg.ml^−1^ hygromycin at 30°C to select for transformants. Colonies were then arrayed on a 96 well plate with 100 µl YPD and grown for 24 hours to induce plasmid loss. The plate is then pinned back onto solid YPD for 24h then replica plated to YPD supplemented with 200 µg.ml^−1^ hygromycin to check for plasmid loss. Allele specific PCR was performed on colonies with loss of plasmid^40^ to distinguish correctly edited allele from wildtype allele. Strains who showed amplification for the edited allele and no amplification for the wildtype allele were phenotyped (4 replicates) on the corresponding condition to measure differences with their wildtype counterparts.

## Acknowledgments

We thank Joshua Bloom and Leonid Kruglyak for insightful discussions, comments on the manuscript as well as for sharing their unpublished manuscript. We thank Maitreya Dunham and the members of the Schacherer laboratory for comments and suggestions. We also thank Gilles Fischer for providing the pAEF5 plasmid. This work was supported by a National Institutes of Health (NIH) grant R01 (GM101091-01) and a European Research Council (ERC) Consolidator grant (772505). T.F. is supported in part by a grant from the Ministère de l’Enseignement Supérieur et de la Recherche and in part by a fellowship from the medical association la Fondation pour la Recherche Médicale. J.S. is a Fellow of the University of Strasbourg Institute for Advanced Study (USIAS) and a member of the Institut Universitaire de France.

