## Supplemental Figures for "Extensive impact of low-frequency variants on the phenotypic landscape at population-scale"

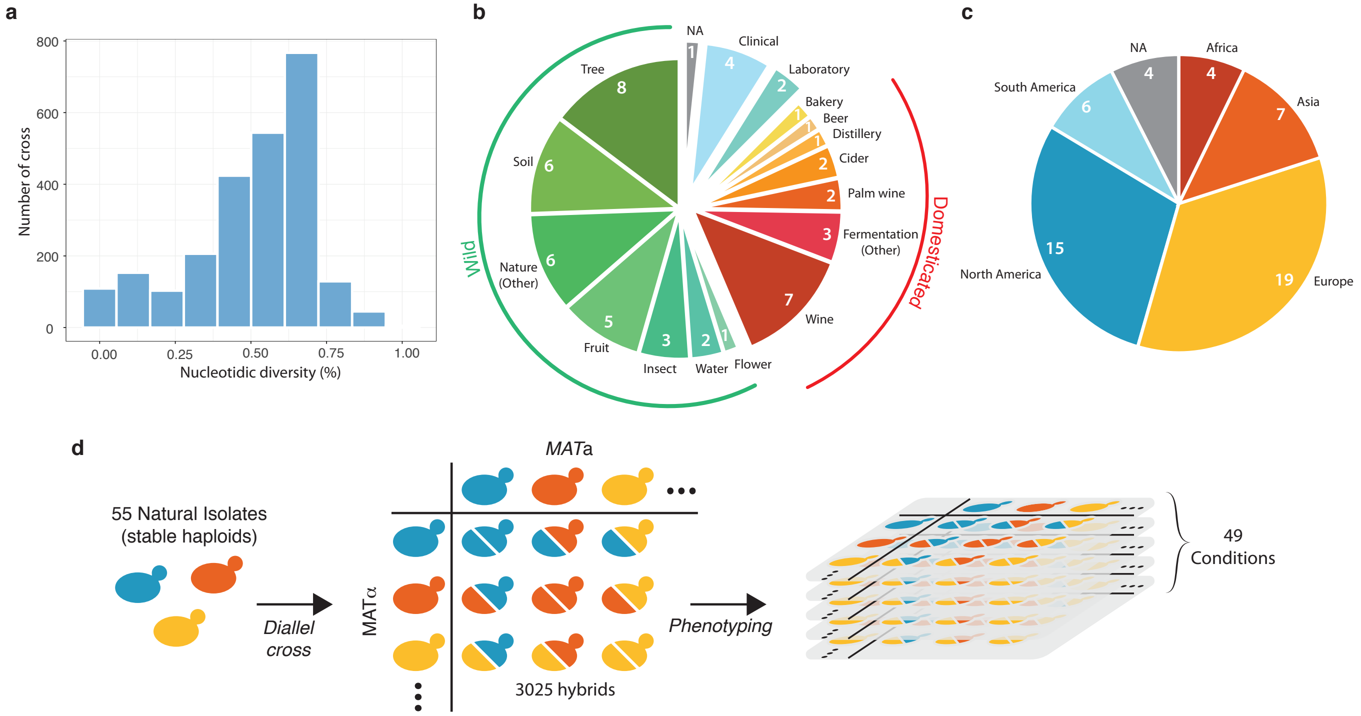


**Supplementary Figure 1 | Diversity of the selected natural isolates and diallel design**
**a.** Pairwise sequence diversity between each pair of parental strains. **b.** Ecological origins of the selected strains. **c.** Geographical origins of the selected strains. **d.** Generation of the diallel hybrid panel. 55 natural isolates available as both mating types as stable haploids were crossed in a pairwise manner to obtain 3,025 hybrids. This panel was then phenotyped on 49 growth conditions impacting various cellular processes.

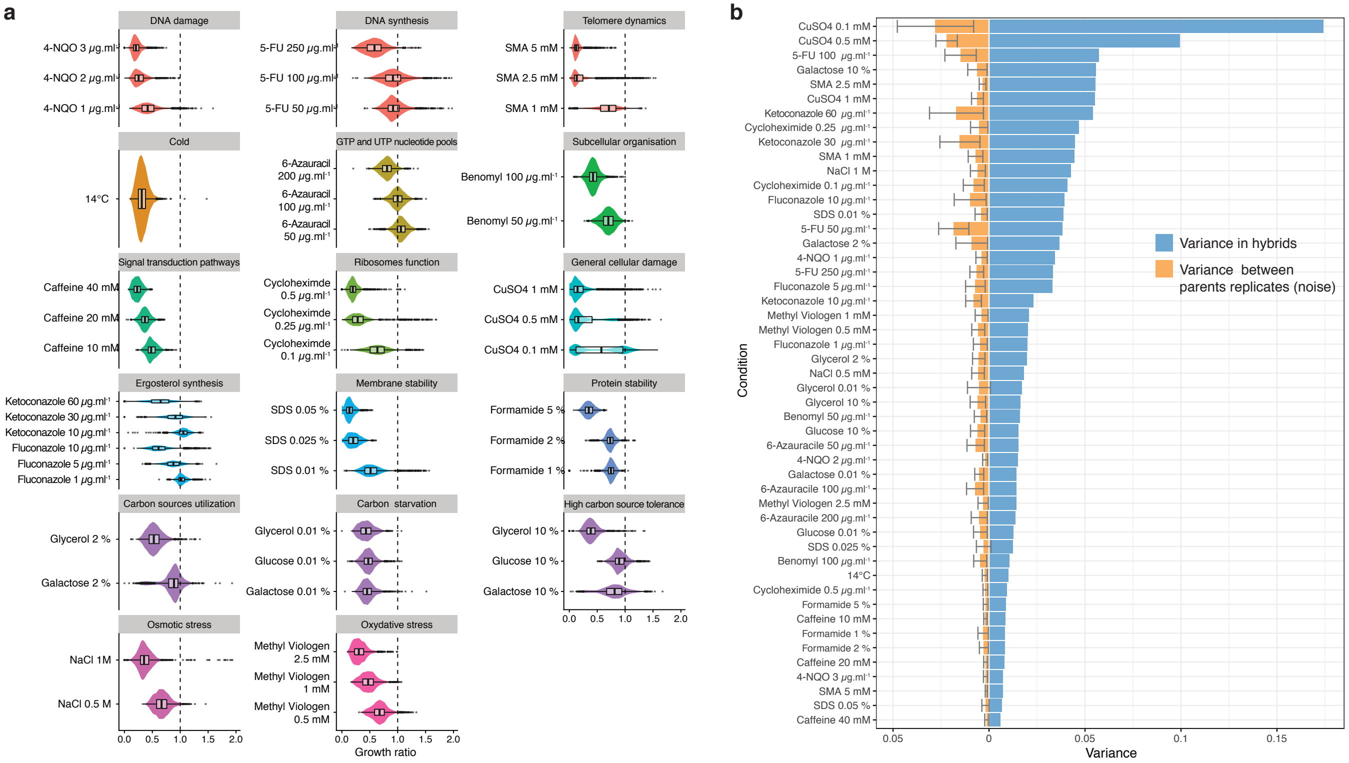

**Supplementary Figure 2 | Phenotypic variance in hybrids**

**a.** Phenotypic distribution for all hybrids in the different growth conditions. Conditions are organized by type of stress in each panel. **b.** Blue bars show the phenotypic variance of the growth ratio for the hybrids in each condition (mean = 0.027). Orange bars represent the variance due to noise between each plate (mean = 0.006). Noise has been measured as the mean variance of every parental replicates across all plates for each condition. Error bars represent interquartile range.

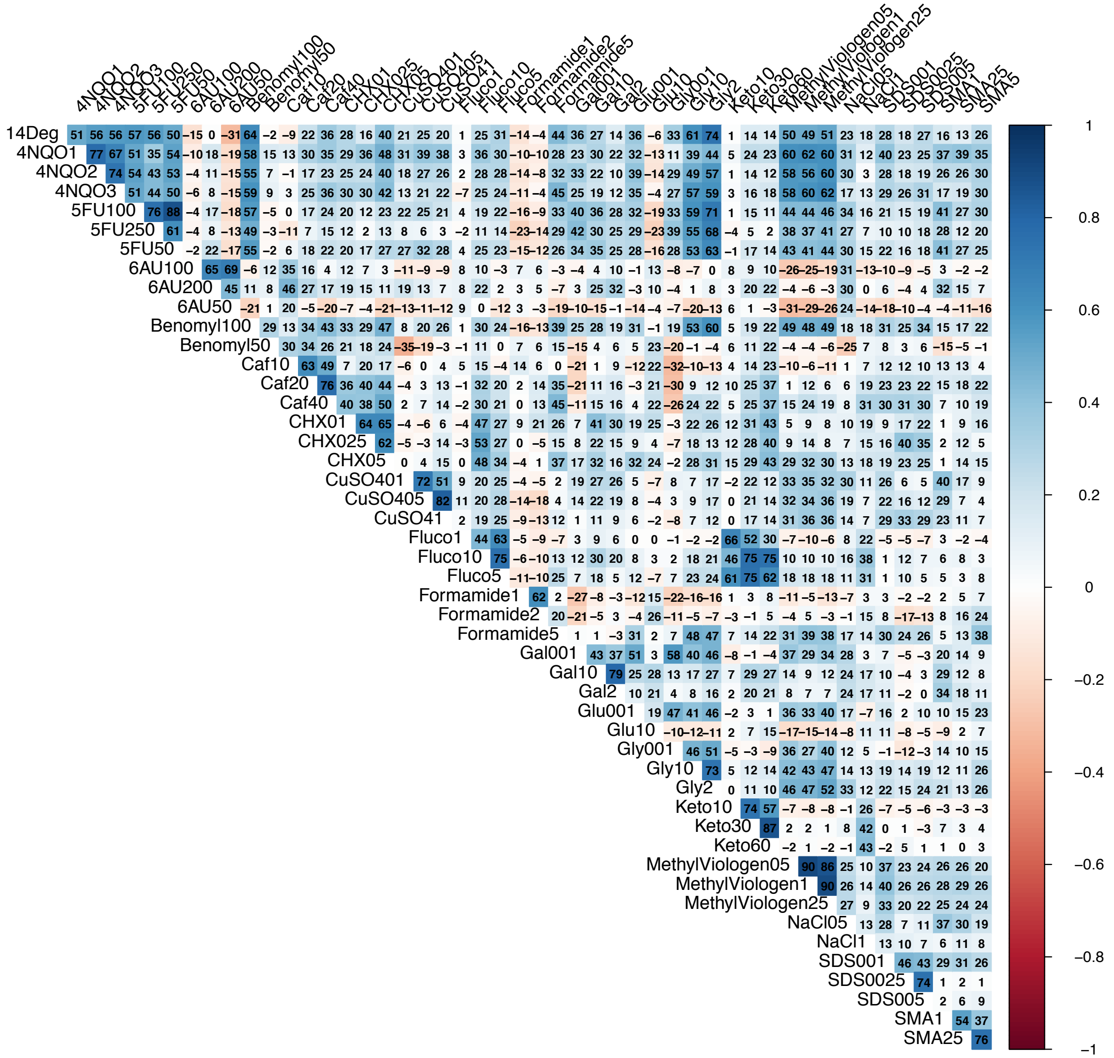

**Supplementary Figure 3 | Correlation between conditions**

Correlogram of all tested growth conditions. Numbers in each cell represent 100 x Pearson's r value.

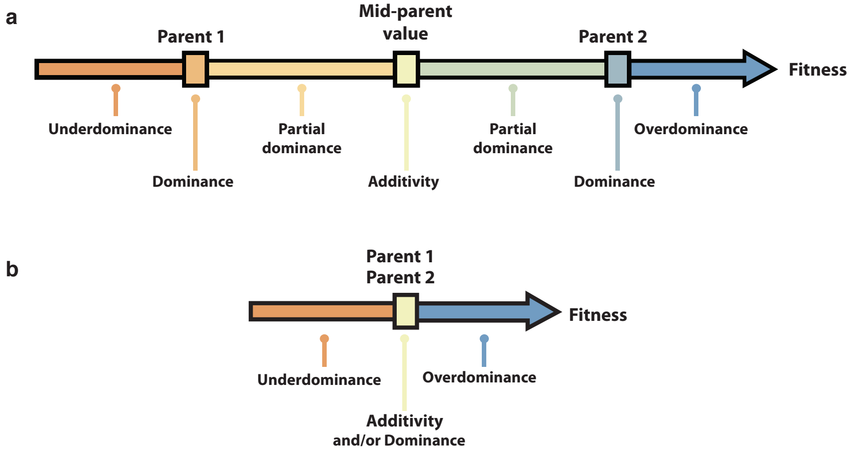


**Supplementary Figure 4 | Classification of inheritance mode**
Representation of the different mode of inheritance depending on the hybrid value: **a.** If a separation can be achieved between parental strains and **b.** If a clear separation cannot be achieved between parental strains.

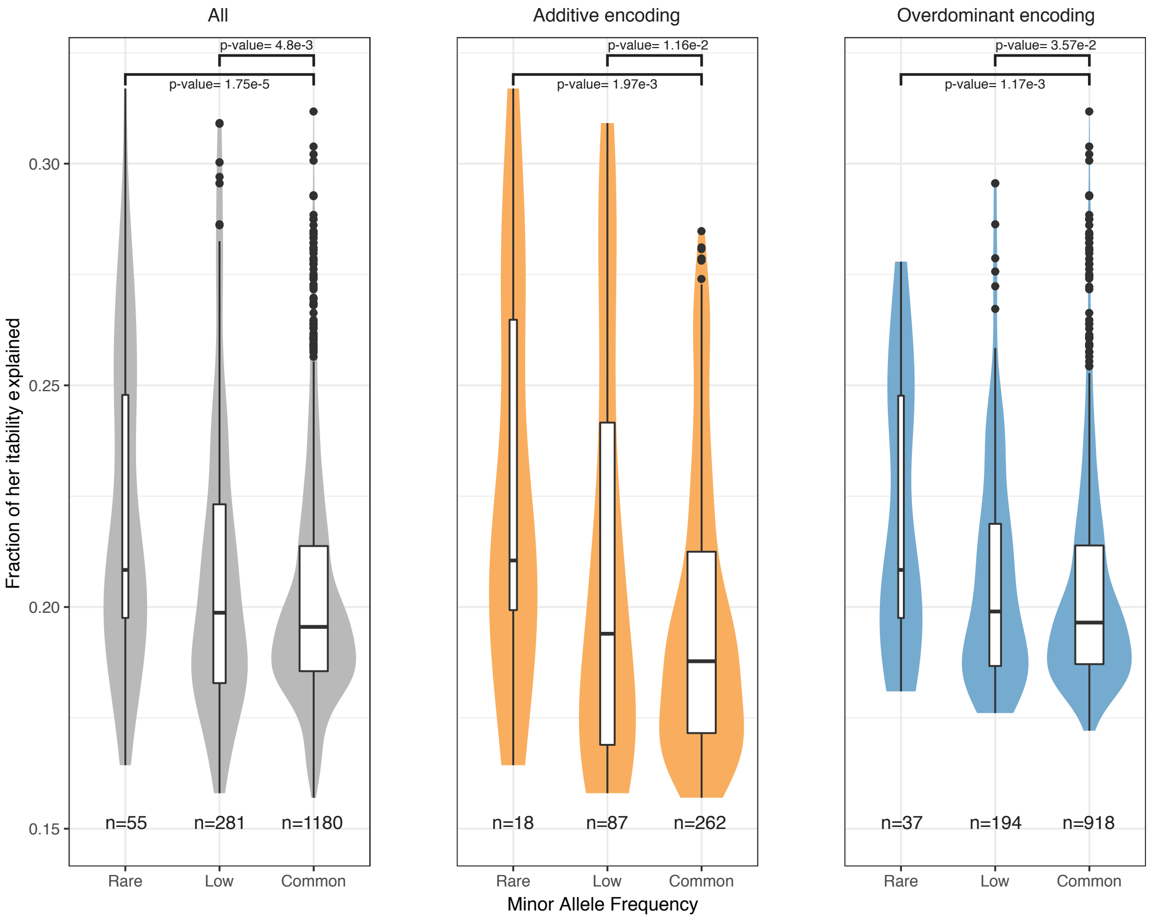


**Supplementary Figure 5 | Variance explained by significantly associated SNPs**
Variance explained for each significantly associated SNPs, for rare (MAF<1%), low-frequency (MAF<5%) and common (MAF>5%) variants for both encoding models (in grey), additive encoding only (in orange) and overdominant encoding (in blue). All p-values are calculated using a two-sided Mann-Whitney-Wilcoxon test.

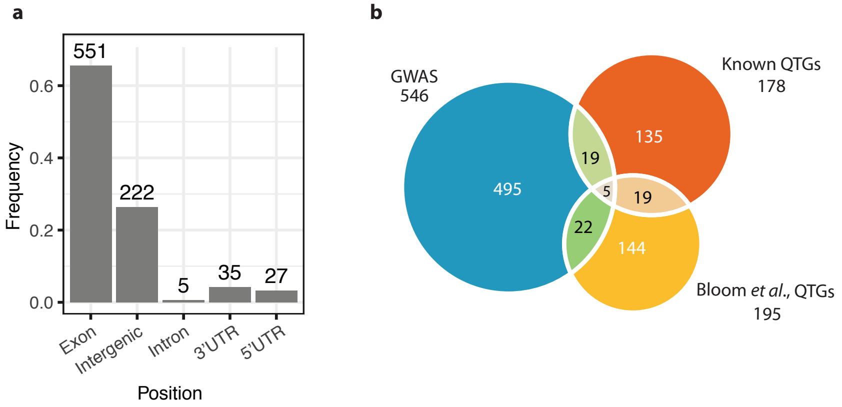


**Supplementary Figure 6 | Significantly associated SNPs position**
**a.** Position of the unique significantly associated SNPs. **b.** Venn diagram comparing the overlap between the 546 unique genes in our dataset with the 178 known QTGs[30] and 195 QTGs recently highlighted[31].

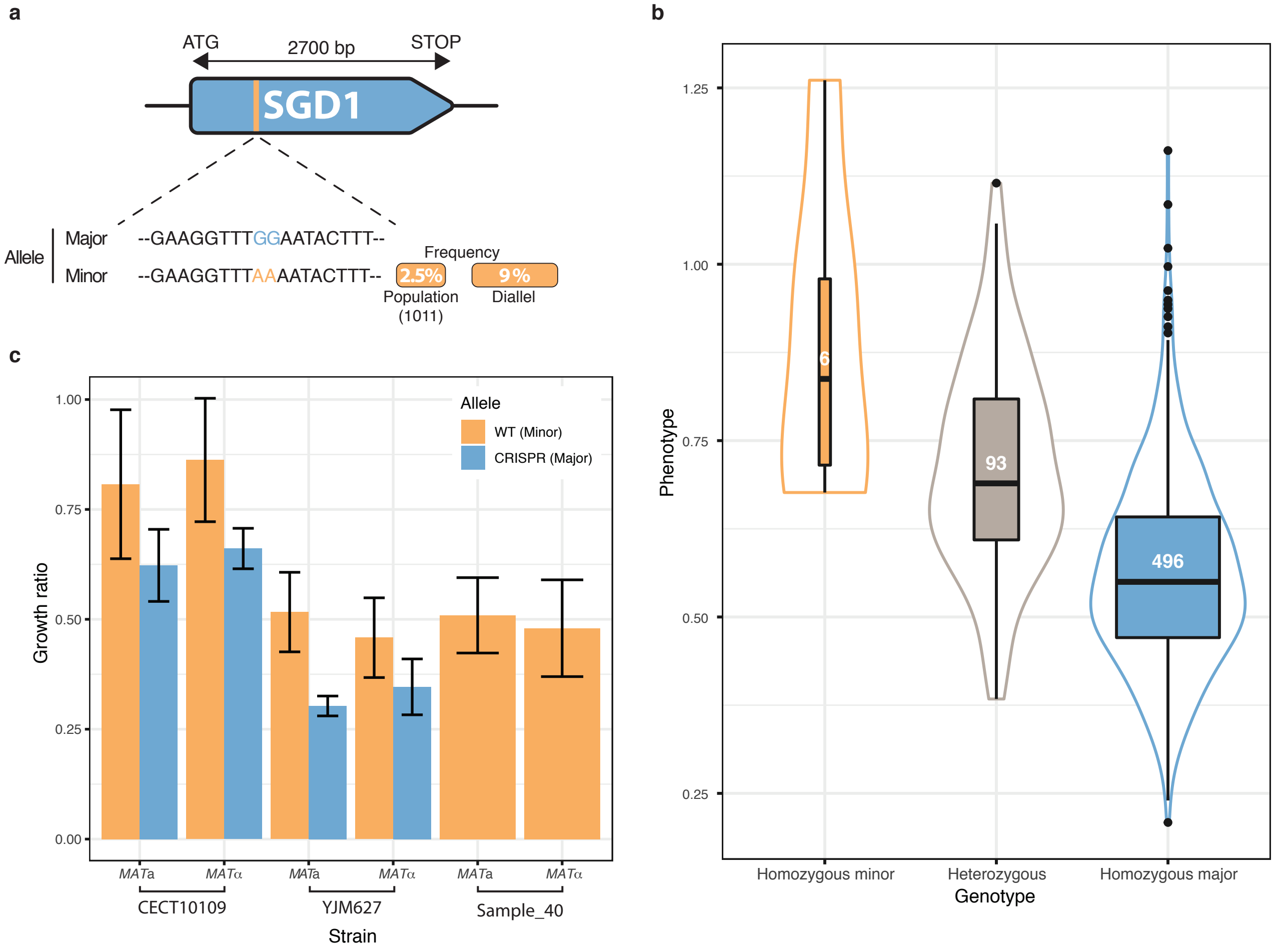


**Supplementary Figure 7 | Low-frequency variant validation in 6-azauracil 100 µg.ml$^{-1}$**

**a.** Schematic representation of SGD1 with the relative position of the detected SNPs. The minor allele is represented in orange with its MAF in the population and in the diallel cross panel b. Boxplot and density plot of the normalized phenotypes for each genotype on 6-azauracile 100 µg.ml-1. Number of observations is displayed inside the boxplots. c. Phenotypic validation after allele replacement of the minor allele with the major allele using CRISPR-Cas9 in the strains carrying the minor allele. Error bars represent median absolute deviation (4 replicates).

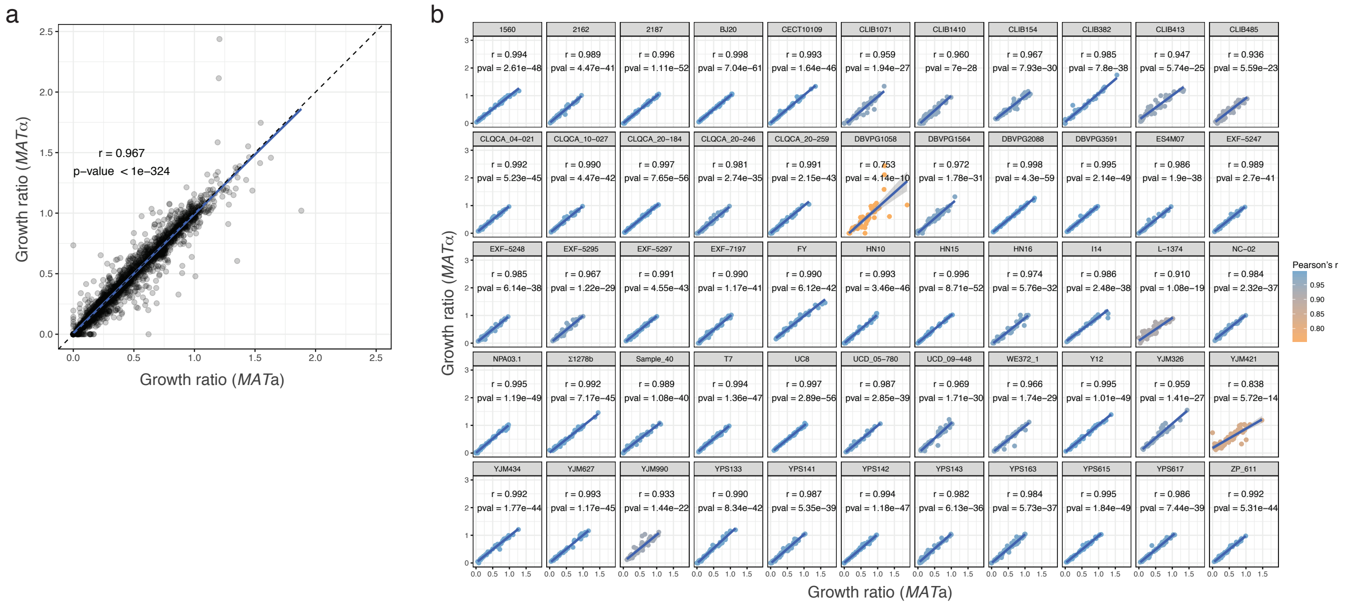


**Supplementary Figure 8 | Phenotypic correlation between *MAT*a and *MAT*α isolates**

**a.** Correlation between growth ratio of different mating types for all parental strains across all conditions. **b.** Correlation between mating types by strain. Pearson's r and corresponding p-values are indicated for each strain.
